# Nuclear upregulation of PI3K p110β correlates with increased rRNA transcription in endometrial cancer cells

**DOI:** 10.1101/2019.12.20.884122

**Authors:** Fatemeh Mazloumi Gavgani, Thomas Karlsson, Ingvild L Tangen, Andrea Papdiné Morovicz, Victoria Smith Arnesen, Diana C. Turcu, Camilla Krakstad, Julie Guillermet-Guibert, Aurélia E Lewis

**Author notes:** equal authorship (Fatemeh Mazloumi Gavgani, Thomas Karlsson). Corresponding author: Aurélia E. Lewis, Department of Biological Sciences, University of Bergen, PO Box 7803, N-5020 Bergen, Norway. The Sars International Centre for Marine Molecular Biology, Bergen 5006, Norway (Fatemeh Mazloumi Gavgani). Department of Biomedicine, University of Bergen (Victoria Smith Arnesen).

## Abstract

Genes encoding for components of the phosphoinositide 3-kinase (PI3K) pathway are frequently mutated in cancer, including inactivating mutations of *PTEN* and activating mutations of *PIK3CA*, encoding the PI3K catalytic subunit p110α. *PIK3CB*, encoding p110β, is rarely mutated, but can contribute to tumourigenesis in some PTEN-deficient tumours. The underlying molecular mechanisms are however poorly understood. By analysing cell lines and annotated clinical samples, we have previously found that p110β is highly expressed in endometrial cancer (EC) cell lines and that *PIK3CB* mRNA levels increase early in primary tumours correlating with lower survival. Selective inhibition of p110α and p110β led to different effects on cell signalling and cell function, p110α activity being correlated to cell survival in *PIK3CA* mutant cells and p110β with cell proliferation in PTEN-deficient cells. To understand the mechanisms governing the differential roles of these isoforms, we assessed their sub-cellular localisation. p110α was cytoplasmic whereas p110β was both cytoplasmic and nuclear with increased levels in both compartments in cancer cells. Immunohistochemistry of p110β in clinically annotated patient tumour sections revealed high nuclear/cytoplasmic staining ratio, which correlated significantly with higher grades. Consistently, the presence of high levels of p110β in the nuclei of EC cells, correlated with high levels of its product phosphatidylinositol 3,4,5-trisphosphate (PtdIns(3,4,5)*P*_3_) in the nucleus. Using immunofluorescence labelling, we observed both p110β and PtdIns(3,4,5)*P*_3_ in the nucleoli of EC cell lines. The production of nucleolar PtdIns(3,4,5)*P*_3_ was dependent upon p110β activity. EC cells with high levels of nuclear PtdIns(3,4,5)*P*_3_ and p110β showed elevated nucleolar activity as assessed by the increase in 47S pre-rRNA transcriptional levels in a p110β-dependent manner. Altogether, these results present a nucleolar role for the PI3K pathway that may contribute to tumour progression in endometrial cancer.

## Introduction

The phosphatidylinositol 3-kinase (PI3K) signalling pathway is frequently hyperactivated in cancer, often due to genetic or epigenetic alterations in several gene members of the pathway [1–3]. Class I PI3Ks consist of heterodimers of catalytic subunits (p110α, β, δ or γ) and adaptor proteins (p85α and its variants, p85β or p55γ) [4] and phosphorylate the 3’-hydroxyl group of the phospholipid phosphatidylinositol 4,5-bisphosphate (PtdIns(4,5)*P*_2_) to generate phosphatidylinositol 3,4,5-trisphosphate (PtdIns(3,4,5)*P*_3_). PtdIns(3,4,5)*P*_3_ binds to effector proteins including the serine/threonine kinases AKT/Protein Kinase B (PKB), 3-phosphoinositide-dependent protein kinase 1 (PDK1) and SIN1 via their phosphoinositide-binding plextrin homology (PH) domain [5]. AKT is activated by phosphorylation on Thr308 and Ser473 by PDK1 and mammalian target of rapamycin complex 2 (mTORC2) respectively [6]. Activated AKT can act at different intracellular sites, where it phosphorylates a myriad of substrates that regulate cell survival, cell proliferation and growth as well as metabolism [7]. The production of PtdIns(3,4,5)*P*_3_ is regulated by phosphatase and tensin homolog deleted on chromosome 10 (PTEN), a lipid phosphatase which dephosphorylates PtdIns(3,4,5)*P*_3_ back to PtdIns(4,5)*P*_2_, thereby opposing PI3K-mediated signalling and hence limiting the potential cancer-promoting effects of class I PI3K activity [8].

p110α and p110β are ubiquitously expressed, share the same enzymatic properties, generate the same lipid product, and initiate the same signalling cascade. Despite these shared features, the two isoforms are both essential in combination for development as individual knockout mice are embryonically lethal, hence suggesting non-redundant functions [9, 10]. Moreover, their mode of activation can be distinct, with p110α carrying out most of receptor tyrosine kinase (RTK)-mediated PI3K signalling and p110β by G-protein coupled receptors (GPCR) as well as RTKs [3, 11–13] through different adaptor proteins [14]. In cancer, the oncogenic properties of p110α are due to activating mutations of its gene *PIK3CA* [15]. In contrast, *PIK3CB*, the gene encoding p110β, is rarely mutated in cancer, with only three reports so far describing activating mutations [16–18]. *PIK3CB* was however shown to be the key isoform mediating tumour growth in PTEN-deficient tumours in particular in breast, prostate and ovarian cancer cells [13, 19–22], possibly due to its ability to promote oncogenic transformation in its wild type state [23]. Furthermore, the importance of p110β in tumourigenesis was recently highlighted in a study by Juric *et al* [24]. This study showed that *PIK3CA* mutant breast cancer cells, which were initially sensitive to p110α specific inhibition, eventually developed resistance with acquired loss of PTEN in metastatic lesions. These cells could however reverse the resistance when p110β inhibition was combined. Regarding their functions, a few studies have reported that the two isoforms can contribute differently to cell survival and proliferation, with p110α playing more of a role in cell survival and p110β in cell cycle progression and DNA replication [25–27]. Another distinguishing feature about these two isoforms is their sub-cellular localisation. Although p110α and β are both found in the cytoplasm and share/compete for similar upstream receptor activation and downstream signalling cascades, p110β harbours a nuclear localisation signal and is found in the nucleoplasm, the chromatin fraction [27, 28] as well as in the nucleolus together with its product PtdIns(3,4,5)*P*_3_ [29]. This would suggest that p110β can orchestrate different processes emanating from the nucleus and explain, at least partly, the pleiotropic aspects of the PI3K pathway.

The PI3K pathway is the signalling pathway most frequently altered in endometrial cancer (EC) with more than 80% of tumours harbouring somatic alteration in at least one gene component of the pathway [30]. This includes high frequency mutations in *PTEN*, *PIK3CA* and *PIK3R1* (encoding p85α) and low frequency in *Akt* and *PIK3R2* (encoding p85β) [31, 32]. Loss of function of the tumour suppressor gene *PTEN*, due to loss of heterozygosity or somatic mutations is the most common event in type I endometrioid EC (EEC) and occurs early in 18-48% of lesions with atypical hyperplasia [33, 34]. *PIK3CA* is the second most frequently mutated in EC with mutations occurring in type I EEC and type II non endometrioid EC serous lesions [32, 34–40]. In addition, mutations in *PTEN* were found to co-exist with those of *PIK3CA* or *PIK3R1*, thereby leading to enhanced activation of the PI3K pathway [35, 39, 41, 42]. *PIK3CA* gene amplification can also account for other mechanisms for PI3K pathway activation and tend to segregate more frequently to aggressive and invasive type II tumours [37, 39, 40, 43]. In contrast to *PIK3CA*, mutation events are rare in *PIK3CB* and account for 2 to 10% in EC according to public data from the Catalog of Somatic Mutations In Cancer (COSMIC, v90) or The Cancer Genome Atlas [37, 40, 44]. In particular, two oncogenic mutations located in its catalytic domain have recently been characterized [17, 45]. *PIK3CB* mRNA levels were found to be elevated in endometrial tumours compared to normal tissue in a few patient samples [46]. In a recent study, we have shown that the p110β protein levels are elevated in EC cell lines and that mRNA levels are increased in grade 1 endometrioid endometrial lesions compared to complex hyperplasias [47]. We have recently reported the presence of p110β and of its product PtdIns(3,4,5)*P*_3_ in the nucleolus of the breast cancer cell line AU565 [29]. In this study, we showed an increase in the nuclear levels of both p110β and PtdIns(3,4,5)*P*_3_ in EC cells. We further showed that high p110β levels correlated with high rRNA transcription, which was partly dependent on p110β activity. These results suggest therefore the involvement of this kinase and its lipid product PtdIns(3,4,5)*P*_3_ in increased nucleolar activity in cancer.

## Results

### p110β is cytoplasmic and nuclear in endometrial cancer cell lines and patient tumours

Previous studies have shown that p110α and p110β are differently localized and that this may contribute to their different cellular function [27–29]. Using cell fractionation and Western immunoblotting, we determined the subcellular localization of PI3K catalytic isoforms, as well as the two regulatory subunits p85α and p85β, in EC cells compared to a non-tumour immortalized endometrial cell line (EM). As shown in Figure 1A, p110α concentrated to the cytoplasmic fraction in all cell lines. p110β expression was low in EM cells detected mostly in the cytoplasmic fraction and with very low levels in nuclear fractions. All cancer cell lines had higher levels of p110β in the cytoplasmic fraction than EM cells. In the nuclear fractions, all cancer cell lines demonstrated high p110β levels except for EFE-184 cells. In the majority of cell lines, p85α was restricted to the cytoplasmic fraction. In contrast, p85β was mostly undetectable in the cytoplasmic fraction in all cells except for MFE-280 cells, but was concentrated to the nuclear fraction, with high levels in EM, KLE, EFE-184 and MFE-280 cells and lower levels in the remaining cells (Figure 1A). To determine if the expression and localization pattern of p110β could also be observed in human tissues, we examined a patient cohort of 727 primary endometrial tumours by immunohistochemistry of tissue microarray (TMA). The level and intensity patterns of p110β were scored separately in the cytoplasm and nucleus. While most patients showed p110β cytoplasmic detection with various degrees of intensity, 23% of all cases showed nuclear staining (Figure 1B). In addition, a significant correlation was observed between high nuclear to cytoplasmic score ratio for p110β and high grade or histological type II (non-endometrioid) endometrial tumour (Figure 1C).

**Figure 1 legend:**
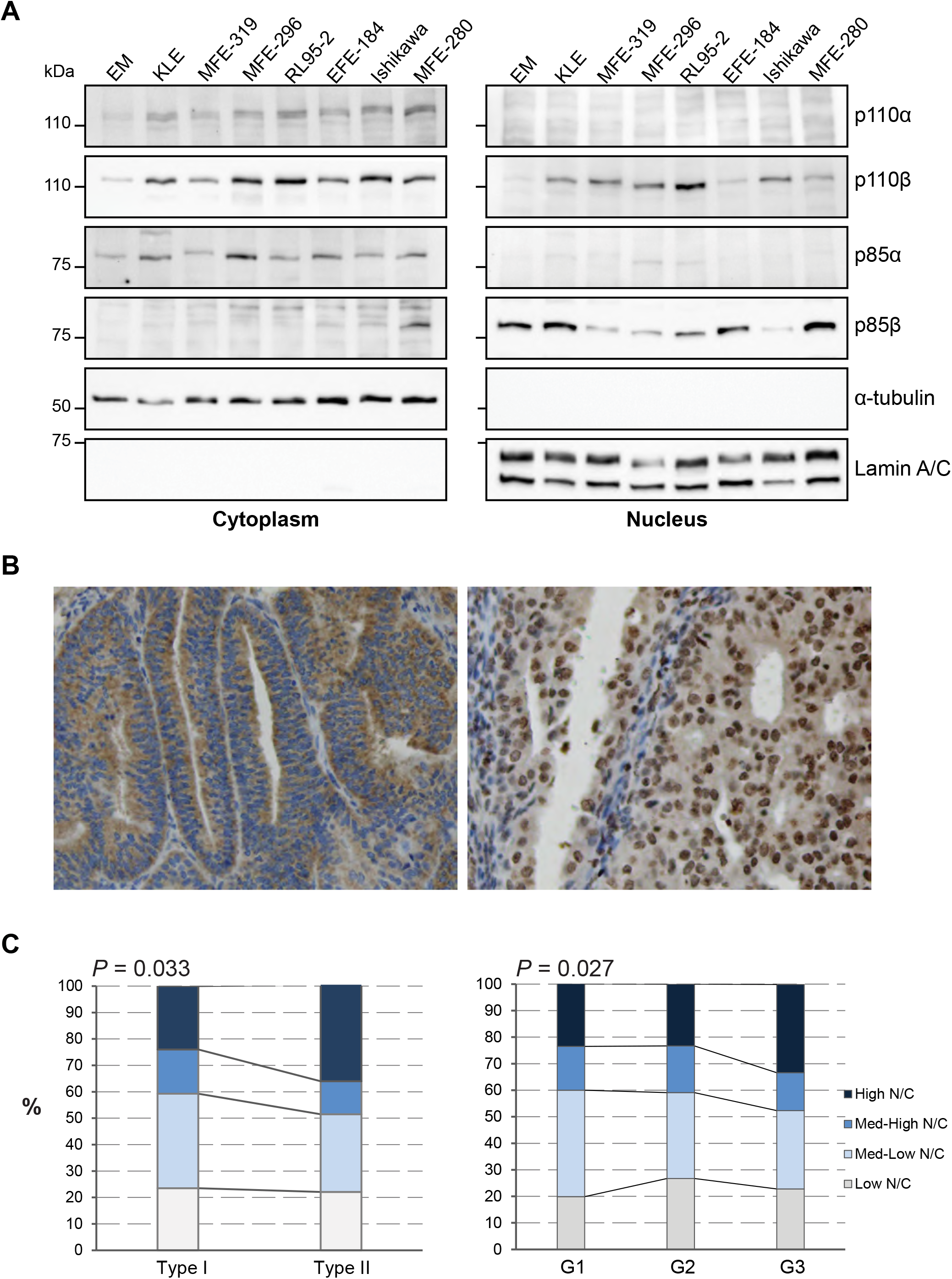
p110β localises to the nucleus in endometrial cancer cells. **(A)** Actively growing cells were fractionated into cytoplasmic and nuclear fractions. Equal protein concentrations were resolved by SDS-PAGE and analysed by Western immunoblotting using the antibodies as indicated. **(B)** Representative histochemistry images of cytoplasmic and nuclear p110β staining in primary endometrial tumours detected with anti-p110β. Magnification x20. **(C)** Quantitative graphs of nuclear (N) to cytoplasmic (C) ratio measured following p110β histochemistry of 714 patient histology samples. G represents the grade of the tumour. The number of samples per grade were as follows: G1= 267, G2= 210, G3= 237.

### The levels of PtdIns(3,4,5)*P*_3_ are increased in nuclei of EC cells in a p110β-dependent manner

We next determined if the presence of p110β in the nucleus correlated with nuclear PI3K pathway activity by first assessing the presence of active p-S473-AKT. As shown in Figure 2A, the cytoplasmic and nuclear levels of p-S473-AKT were low in EM, KLE, EFE-184 and MFE-280 cells, while high levels were observed in PTEN-deficient cells, MFE-296, MFE-319, RL95-2 and Ishikawa cells, consistent with our previous study using total cell extracts in the same cells [47]. Interestingly, high nuclear p-S473-AKT levels were inversely correlated with low levels of p85β (Figure 1A). Furthermore, we determined the nuclear level of PtdIns(3,4,5)*P*_3_ of all cells examined following nuclear isolation, lipid extraction, and detection with GST-GRP1-PH, a PtdIns(3,4,5)*P*_3_ specific probe ([48] Figure 2B and Supplementary Figure S1 showing the specificity of the probe). The purity of the fractionation was verified by Western immunoblotting using markers for the cytoplasm, nucleus and endoplasmic reticulum (Supplementary Figure S2). PtdIns(3,4,5)*P*_3_ levels were high in most cancer cells and highest in RL95-2 cells compared to EM cells (Figure 2C). To test if p110β is responsible for the synthesis of nuclear PtdIns(3,4,5)*P*_3_, we treated the PTEN-deficient cell line RL95-2 with TGX-221, a p110β selective inhibitor. Treatment for 3 days reduced the levels of nuclear PtdIns(3,4,5)*P*_3_ (Figure 2D) as well as nuclear p-S473-AKT (Figure 2E). However, the levels of total AKT were increased in the cytoplasm while it was decreased in the nucleus following p110β inhibition. The decrease in nuclear p-AKT may hence be due to loss of translocation of active pAKT from the cytoplasm. Knockdown of p110β also led to a decrease in the nuclear levels of PtdIns(3,4,5)*P*_3_ and p-S473-AKT (Figure 2F-G).

**Figure 2 legend:**
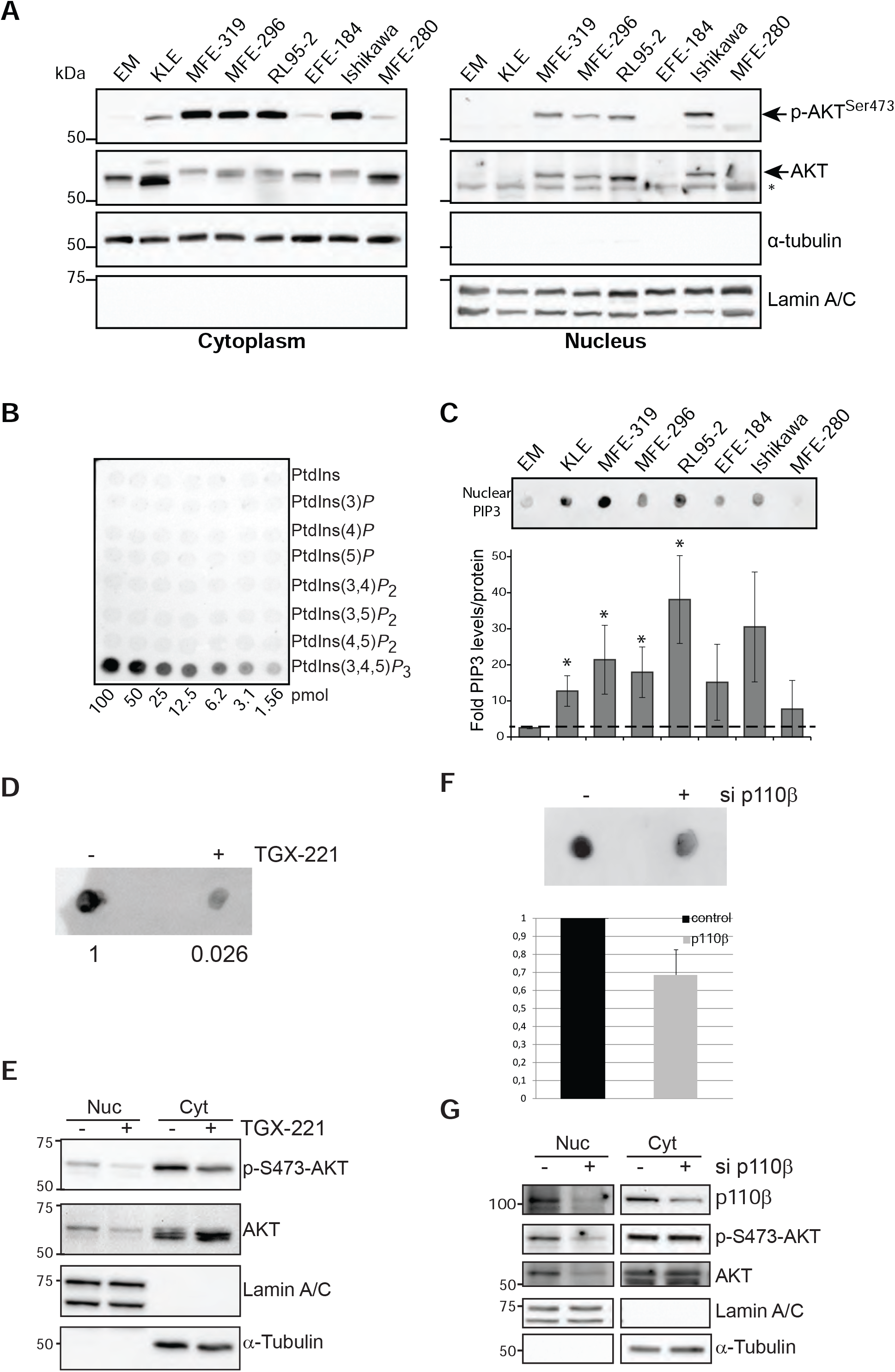
Nuclear PtdIns(3,4,5)*P*_3_ levels are elevated in endometrial cancer cells. **(A)** Actively growing cells were fractionated into cytoplasmic and nuclear fractions. Equal protein concentrations were resolved by SDS-PAGE and analysed by Western immunoblotting using the antibodies as indicated. **(B)** PIP array spotted with 1.56 to 100 picoM of each of the seven PPIn species incubated with GST-GRP1-PH and an anti-GST-HRP conjugated antibody. **(C)** PtdIns(3,4,5)*P*_3_ (PIP3) detection from nuclear acidic lipids extracted from actively growing cells, by overlay assay with GST-GRP1-PH domain and anti-GST-HRP conjugated antibody (upper panel). PIP3 signal/mg nuclear protein were calculated and expressed as folds compared to EM values (lower panel–graph. n=3, * p< 0.05 T-test). **(D)** PIP3 detection by overlay assay with GST-GRP1-PH domain and anti-GST-HRP conjugated antibody from nuclear acidic lipids extracted from RL95-2 cells treated with or without 10 μM TGX-221 for three days. **(E)** Western immunoblotting of cytoplasmic (cyt) and nuclear (nuc) fractions from RL95-2 cells treated with or without 10 μM TGX-221 for three days. **(F)** PIP3 detection by overlay assay with GST-GRP1-PH domain and anti-GST-HRP conjugated antibody from nuclear acidic lipids extracted from RL95-2 cells treated with 200 nm control or p110β siRNA. **(G)** Western immunoblotting of cytoplasmic (cyt) and nuclear (nuc) fractions from RL95-2 cells treated with 200 nM control or p110β siRNA.

### p110β and PtdIns(3,4,5)*P*_3_ are nucleolar in EC cells

Consistent with our previous study [29], we found that p110β was localized in the cytoplasm, the nucleoplasm and strongly in nucleoli together with the nucleolar protein nucleophosmin in three cell lines (Figure 3A) with high nuclear p110β. The specificity of the anti-p110β antibody was validated by knockdown using siRNA (Figure 3B). Nucleolar fractionation of RL95-2 cells confirmed the presence of p110β in the same compartments by Western immunoblotting (Figure 3C). α-Tubulin, lamin A/C and fibrillarin were used as cytoplasmic, nuclear and nucleolar markers, respectively to validate the fractionation procedure. Lamin A/C was found in both the nucleoplasmic and nucleolar compartments as previously reported [49]. p110β was also detected with PtdIns(3,4,5)*P*_3_ both in the nucleoplasmic and nucleolar compartments of RL95-2 (Figure 3D) and MFE-319 cells (Supplementary Figure S3). The specificity of the anti-PtdIns(3,4,5)*P*_3_ antibody was validated by lipid overlay assays using PIP arrays and by competition with free lipids by immunofluorescence (Supplementary Figure S4). Hence, pre-incubation of the antibody with different PPIns showed that nucleoplasmic and nucleolar staining was abolished by the presence of PtdIns(3,4,5)*P*_3_ but not by PtdIns3*P* and PtdIns(3,4)*P*_2_. Furthermore, AKT was found to colocalise with the nucleolar pool of p110β and its active p-S473 form with the nucleolar protein nucleophosmin as discrete foci (Figure 3E).

**Figure 3 legend:**
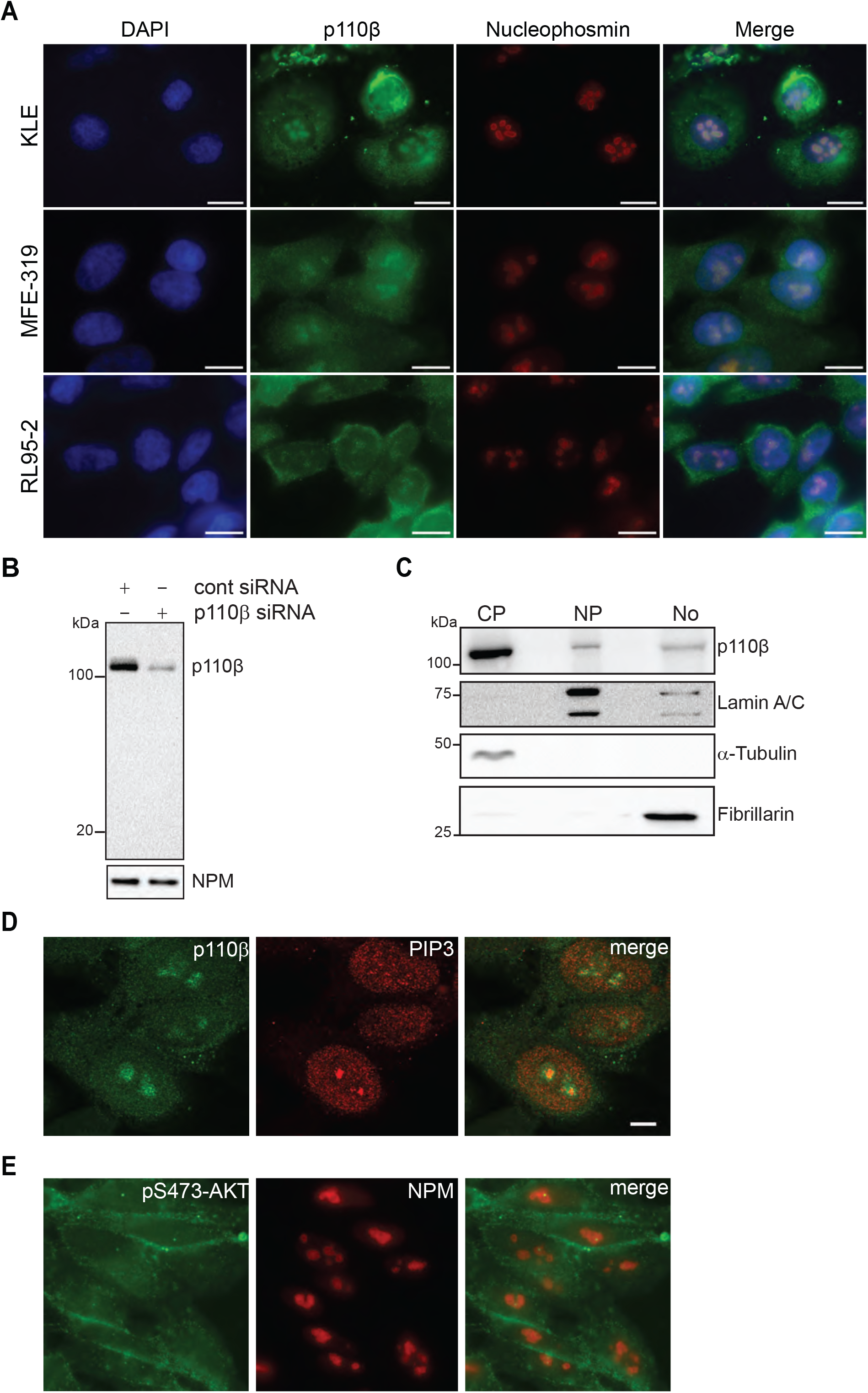
p110β and PtdIns(3,4,5)*P*_3_ are nucleolar. **(A)** Co-immunostaining of p110β using the abcam 151549 anti-p110β antibody and nucleophosmin in actively growing KLE, MFE-319 and RL95-2 cells and imaged by epifluorescence microscopy. Scale bar 10 μm (x100). **(B)** Western immunoblotting of cell extracts from KLE cells treated with 100 nM of control or p110β siRNA, using the abcam 151549 anti-p110β antibody. **(C)** Sub-cellular fractionation of RL95-2 cells and Western immunoblotting of equal protein amounts from each fraction **(D-E)** Confocal microscopy of actively growing RL95-2 cells co-stained with the indicated antibodies (Scale bar 5 μm).

### The nucleolar pool of PtdIns(3,4,5)*P*_3_ is partly dependent upon the activity of p110β

The main function of the nucleolus is to synthesise ribosomes, requiring rRNA transcription, processing and assembly with ribosomal proteins [50]. rRNA transcription oscillates during the cell cycle, as it is lowest during mitosis and is re-activated in G1 phase with highest activity thereafter in S and G2 phases [51]. In parallel, p110β is activated during G1 in the nucleus and contributes to G1 to S phase transition [26, 27] and we have found that p110β, localises dynamically in nucleoli when they start to reform as the cells exit mitosis (Supplementary Figure S5). To determine if the pool of PtdIns(3,4,5)*P*_3_ present in nucleoli is produced due to the kinase activity of p110β, we compared the nucleolar appearance of PtdIns(3,4,5)*P*_3_ in WT and p110β kinase inactive mouse embryonic fibroblast (MEF) cells during the reformation of nucleoli as cells exit mitosis (Figure 4A). We combined nocodazol treatment and mitotic shake-off to synchronize and enrich for mitotic cells. After plating the collected mitotic cells on coverslips, the cells were left to recover for 4-5 h before they were fixed and labelled with a GFP-GRP1-PH probe and immunostained with anti-nucleophosmin. In line with our results in HeLa cells (Supplementary Figure S5), PtdIns(3,4,5)*P*_3_ was detected together with nucleophosmin in the p110β WT MEFs. In contrast, the p110β kinase inactive MEFs demonstrated a substantial decrease in PtdIns(3,4,5)*P*_3_ nucleolar staining.

**Figure 4 legend:**
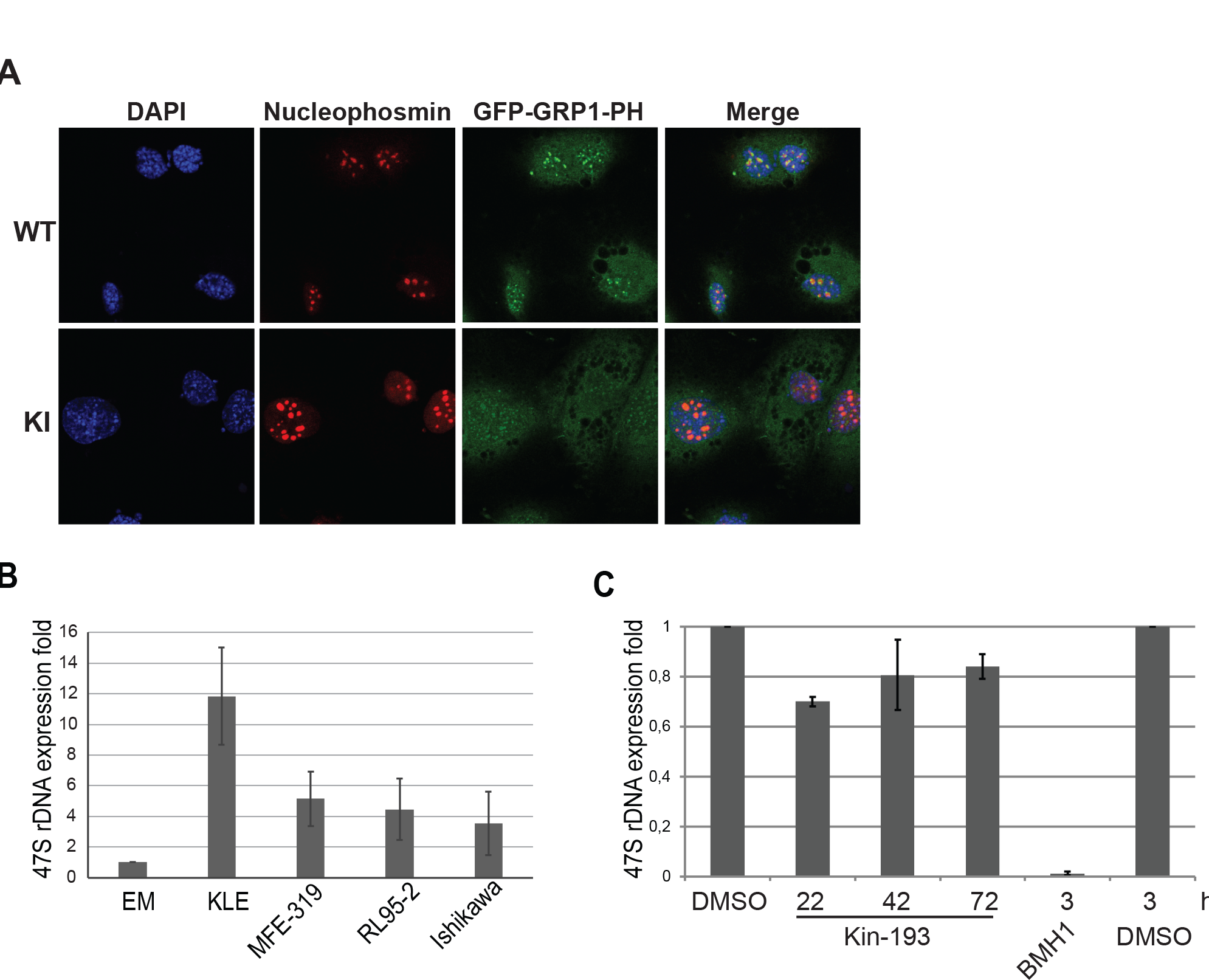
p110β is active in the nucleolus and contributes to 47S pre-rRNA transcription. (A)MEFs p110β WT and Kinase inactive (KI) were treated for 16 h with 50 ng/ml nocodazole, collected after mitotic shake-off and replated. Cells were fixed after 4 h and immunofluorescence staining was performed as indicated followed by confocal microscopy. (B)Relative 47S pre-rRNA to β*-ACTIN* mRNA expression in RL95-2, MFE 319, Ishikawa and EM cells. The expressions of all samples were normalized to EM cells. **(C)** Relative 47S pre-rRNA to β*-ACTIN* mRNA expression in RL95-2 treated in the presence of DMSO or the p110β inhibitor KIN-193 (10 μM) for 22, 42 and 72 h, or the BMH21 (1 μM) for 3 h.

### High nucleolar p110β levels correlates with high 47S rRNA transcription

Elevated levels of nucleolar activity have been correlated to an increased risk of cancer development [52]. We next examined whether EC cells with high levels of p110β and PtdIns(3,4,5)*P*_3_ in the nucleus had high level of 47S rRNA transcription compared to EM cells by quantitative RT-PCR. Consistently, all EC cell lines examined demonstrated higher levels compared to EM cells, with KLE cells, showing the highest levels (Figure 4B). Using a selective inhibitor of p110β, we next tested if p110β is implicated in 47S transcription. A 30% decrease was observed following 22 h of treatment respectively while RNA Pol I inhibition almost completely blocked transcription (Figure 4C).

## Discussion

Cellular compartmentalisation provides an additional important mode of regulation for signalling cascades to achieve specificity and to precisely coordinate cellular outputs. The PI3K pathway has been extensively studied from a cytoplasmic perspective. However, a few studies have detailed the distinct intracellular localisation of PI3K enzymes. The PI3K p110α is found to be restricted to the cytoplasm while p110β is present both in the cytoplasm and the nucleus, and in particular in the nucleoplasm, the chromatin fraction and the nucleolus [27–29]. The compartmentalisation of these enzymes is likely to impact signalling networks and to mediate different cell functions, hence accounting for the pleiotropic effects attributed to PI3K signalling. Although the PI3K signalling pathway is pivotal in cancer, the impact of the subcellular localisation of PI3K in processes attributed to tumourigenesis is still poorly understood. Our findings demonstrate for the first time that p110α and p110β are differently compartmentalised in EC cells. Consistent with previous studies in other cell types [27, 28], p110α is cytoplasmic while p110β is both cytoplasmic and nuclear. This would suggest that, in the cytoplasm, p110α and p110β isoforms can share some of the functions attributed to PI3K signalling operating perhaps due to their reported cross-activation [53]. In cancer cells, the presence of genetic mutations affecting *PIK3CA* or *PTEN* would influence PtdIns(3,4,5)*P*_3_-mediated downstream functions induced by p110α and p110β respectively in this compartment. Concerning the nucleus, we found that the levels of p110β are high in the nucleus of EC cell lines compared to EM cells. In clinically annotated tumour samples, we showed a correlation between the nuclear p110β levels and endometrial cancer progression as tumours with higher grade presented high p110β nuclear to cytoplasmic ratio. These results would indicate the importance of the levels of this isoform combined with its compartmentalisation status. At least, this would be consistent with the fact that the overexpression of p110β has previously been shown to lead to cell transformation in its wild type state [23]. However, this is the first study showing the potential importance of an increase in the nuclear compartmentalisation of p110β during disease progression in clinical samples.

Furthermore, our studies demonstrate that EC cells, not only have high nuclear levels of p110β, but also elevated levels of PtdIns(3,4,5)*P*_3_, its lipid product as well as the active form of the oncoprotein AKT, p-S473-Akt, the key signalling effector of PtdIns(3,4,5)*P*_3_. Here, we demonstrate that upon p110β inhibition, the levels of p-S473-Akt were decreased in EC nuclei. The nuclear PtdIns(3,4,5)*P*_3_ levels were also reduced in these cells following p110β inhibition, which suggests that the nuclear pool of PtdIns(3,4,5)*P*_3_ is, at least partly, the product of the kinase activity of p110β. Another pool could be dependent upon the activity of inositol polyphosphate kinase (IPMK). IPMK was initially discovered as an inositol(1,4,5)triphosphate kinase but was thereafter reported to also act on PPIn to generate PtdIns(3,4,5)*P*_3_ [54, 55]. The existence of a molecular link within the nucleus between PtdIns(3,4,5)*P*_3_ and Akt is however not clear from this study. Additional mechanisms of regulation required for the activation of Akt were not explored. These may include the PtdIns(3,4,5)*P*_3_-dependent activation of PDK1 and mTORC2, known to be critical for the phosphorylation and full activation of Akt. This may be plausible as they have been detected in the nucleus in previous studies [56–59].

A clear relationship between elevated nucleolar activity and increased risk of cancer has been shown [52]. Thus, nucleolar processes need to be tightly regulated with high fidelity to ensure appropriate cell growth and proliferation in response to external signals. One potential molecular link regulating these processes is the PI3K pathway. Previous studies have shown that the transcription and processing of the pre rRNA is stimulated in a PI3K and mTOR-dependent manner [60–62]. However, the responsible PI3K isoform was not identified in those studies. Interestingly, the PtdIns(3,4,5)*P*_3_ effector protein nucleophosmin [63] as well as mTORC1 are known to localize to the nucleolus, where they were shown to regulate nucleolar function [62, 64–66]. Nuclear Akt has also been shown to regulate rRNA transcription by activating the TIF-I transcription factor [67]. This would correlate well with our findings detecting pAKT in the nucleolus. These studies would suggest that the key members of the PI3K pathway are present in the nucleolus but how PI3K, Akt and mTOR are activated in a nucleolar context, i.e. in a membrane-less environment is however unclear. In this study, we showed that both PtdIns(3,4,5)*P*_3_ and p110β were localised in the nucleolar compartment, raising the possibility of p110β acting as a regulator of nucleolar functions in a kinase-dependent manner. Preliminary data showed the detection of the PtdIns(3,4,5)*P*_3_ C38:4 molecular species from isolated nucleoli of RL95-2 cells by LC-MS/MS analyses (data not shown), which is consistent with the reported most common chemical form of fatty acyl chains for polyphosphoinositides (PPIn) [68–70]. Immunofluorescent staining indicated also of the presence of both total Akt and phosphorylated Akt in nucleoli, which can suggest a local activation of this protein by PtdIns(3,4,5)*P*_3_ present in nucleoli. Again, this would need to be explored further, in particular in terms of the biophysical state of PtdIns(3,4,5)*P*_3_ in a membrane-less environment.

Our findings demonstrate that the RL95-2 endometrial cancer cells, high in nuclear PtdIns(3,4,5)*P*_3_ and p110β levels, have significantly increased pre-rRNA transcription. The proliferation of these cells was shown to be at least partly dependent upon the activity of p110β [47]. This would suggest that an increase in ribosome production will help increase cell proliferation and subsequently cancer progression. Nucleolar p110β would hence provide a mode of regulation of ribosome synthesis necessary for protein synthesis and ultimately cell division. p110β can therefore potentially increase tumour progression in EC cells by producing the nucleolar pool of PtdIns(3,4,5)*P*_3_ and thereby increasing the biogenesis of ribosomes required for tumour growth. However, the exact molecular mechanisms by which PtdIns(3,4,5)*P*_3_ or p110β can influence nucleolar function remains to be further explored.

## Materials and methods

### Reagents

Antibodies used in Western immunoblotting and immunostaining are listed in Supplementary Table S1. The selective p110β inhibitors TGX-221 (Cayman chemicals) and KIN-193 (aka AZD6482, from Selleckchem) were dissolved in DMSO at 10 mM and stored at −80°C. The RNA polymerase I inhibitor, BMH21 (from Selleckchem) was dissolved in DMSO at 1 mM. *PIK3CB* (sc-37269) and control non-targeting (sc-37269C) siRNA and transfection reagent were from Santa Cruz. The pGEX-4T1-EGFP-GRP1-PH plasmid was from Dr Julien Viaud (INSERM U.1048, Toulouse, France).

### Protein expression and purification

The pGEX-4T1-EGFP-GRP1-PH plasmid was transformed into *E. coli* BL21-RIL DE3. The bacteria were grown at 37°C and further induced with 0.5 mM isopropyl-β-D-thiogalactopyranoside overnight at 18°C. Bacterial pellets were lysed in 50 mM Tris pH 7.5, 150 mM NaCl, 1% Triton-X100, 5 mM DTT, 10 % glycerol containing protease inhibitor cocktail and 0.5 mg/ml lysozyme, for 30 min on ice. Following sonication and centrifugation, GST-EGFP-GRP1-PH was purified with glutathione-agarose 4B beads, analysed by SDS-PAGE and Coomassie staining.

### Cell lines and cell culture conditions

Cancer cell lines were obtained from ATCC (KLE, RL95-2), DSMZ Germany (MFE-296, MFE-319, EFE-184 and MFE-280) and Sigma-Aldrich (Ishikawa). EM-E6/E7-hTERT (EM), a non-transformed endometrial cell line isolated from glandular endometrial tissue and immortalized with E6/E7 and human TERT [71, 72], was a gift from Professor PM Pollock (University of Queensland, Australia). All cells were authenticated by short tandem repeat DNA profiling (IdentiCell Service, Dept. Molecular Medicine, Aarhus University Hospital, Denmark for all cancer cell lines and MD Anderson Cancer Center, USA for EM cells), as previously described [47]. p110β^D931A/D931A^ kinase inactive and p110β^WT/WT^ MEFs immortalised by Shp53 were from Dr Julie Guillermet-Guibert (Université Toulouse III-Paul Sabatier, Toulouse, France). All EC cells, HeLa and MEFs were cultured in Dulbecco’s modified Eagle’s medium (DMEM) supplemented with 10% foetal bovine serum (FBS) and antibiotics (100 IU/ml penicillin and 100 μg/ml streptomycin). EM cells were cultured in DMEM/Ham’s F12 supplemented with Insulin-Transferrin-Selenium, 10% FBS and antibiotics and changed to DMEM containing 10% FBS and antibiotics 24 h before harvest. Cells were harvested when they reached a maximum of 80% confluence.

### Cell synchronization

Cells grown up to 70% confluency were treated with 50 ng/mL of nocodazole for 16 h. The mitotic cells were collected by mechanical shake-off (repeated twice). The cells were pelleted by centrifugation at 70 g for 5 min. After washing the pellet twice with 10 ml of growth medium the cell pellet was plated on coverslips covered with poly-L-Lysine. The cells were collected at different time points after re-plating.

### Whole cell extracts, subcellular fractionation and Western immunoblotting

Whole cell extracts were prepared in radioimmunoprecipitation assay (RIPA) lysis buffer (50 mM Tris pH 8.0, 0.5% deoxycholic acid, 150 mM NaCl, 1% NP-40, 0.1% SDS) supplemented with 5 mM NaF, 2 mM Na_3_VO_4_ and 1x Sigma Protease Inhibitor Cocktail. Nuclear fractionation was carried out according to O’Caroll *et al.* [73] and nuclear pellets were lysed in RIPA buffer. RL95-2 cells required an additional syringing step of the nuclear pellet resuspended in wash buffer (10 mM Tris-HCl pH 7.5 and 2 mM MgCl_2_) to avoid cytoplasmic contamination. The nucleolar fractionation was performed according to the protocol described in Lam *et al* with minor changes [74]. In brief, cells were grown in 10 × 15 cm dishes up to 70% confluency. Fresh medium was added to the cells 1 h prior to the fractionation. Cells were trypsinized and washed three times with cold PBS. The cell pellet was collected by centrifugation and re-suspend in 5 ml of buffer A containing 10 mM HEPES pH 7.9, 1.5 mM MgCl_2_, 10 mM KCl, 1% Igepal and protease inhibitor cocktail. After 5 min of incubation on ice, the cells were syringed 16 times using a 23-gauge needle. After centrifugation at 200 g for 5 min at 4°C the supernatant was collected as the cytosolic fraction and the nuclear pellet was re-suspended in 3 ml of buffer S1 (0.25 M sucrose, 10 mM MgCl_2_ and protease inhibitor cocktail). The suspension was layered over 3 ml of buffer S2 (0.35 M sucrose, 0.5 mM MgCl_2_ and protease inhibitor cocktail) and centrifugation was performed at 1400 g for 5 min at 4°C. The pellet was then re-suspended in 3 ml of buffer S2 and sonicated (7 times: 10 sec on/10sec off) on ice. The lysate was then layered over 3 ml of buffer S3 (0.88 M Sucrose, 0.5 mM MgCl_2_ and protease inhibitor cocktail) and centrifugation was performed at 3000 g for 10 min at 4°C. The top layer was collected as the nucleoplasmic fraction and the pellet which contained the nucleoli was washed once with 500 μl of S2 and centrifuged. Nucleolar pellets were resuspended in RIPA buffer. Equal amount of proteins (40-50 μg) were resolved on denaturing SDS-polyacrylamide gels, immunoblotted on to 0.45 μm thick nitrocellulose membranes and detected by enhanced chemiluminescence using the SuperSignal West Pico Chemiluminescent Substrate (Thermo-Fisher) and visualized with a BioRad ChemiDocTM Xrs+.

### RNA extraction and reverse transcription

Cells were pelleted, washed two times with PBS and resuspended in 1 ml TriReagent (Sigma) and incubated at room temperature for 5 min. 200 μl of chloroform was added, mixed vigorously, incubated at room temperature for 1 min and centrifuged at 12000 g and 4°C for 15 min. Phenol-chloroform-isoamyl alcohol mixture (Sigma) was added (500 μl) to the upper phase, mixed, incubated at room temperature for 2 min and centrifuged at 12000 g and 4°C for 10 min. Chloroform (500 μl) was added to the upper phase, mixed, incubated at room temperature for 1 min and centrifuged at 12000 g and 4°C for 10 min. 20 μg of RNA grade glycogen (Thermo Fisher Scientific) and 500 μl isopropanol were added to the upper phase, mixed and incubated at room temperature for 20 min before centrifuging at 13000 g and 4°C for 20 min. The pellet was resuspended in 1 ml of ice cold ethanol (70%) and centrifuged (at 8000 g and 4°C) for 5 min. The extracted RNA was dissolved in RNAse-free water for RT-qPCR analysis. cDNA was generated from 1 μg of RNA using random primers according to the protocol from the High-Capacity cDNA Reverse Transcription Kit (Thermo Fisher scientific) with RNase Inhibitor.

### Real time qPCR

Real-time qPCR was performed from three biological replicates in triplicates on the Roche Light Cycler 480 using PowerUp™ SYBR™ Green Master Mix (Thermo Fisher scientific or Life Sciences). The reaction mix contained 10 μM of each primer and 1 ng of cDNA diluted. The primers used for the target human 47S-rRNA allow the amplification of the sequence (+302 and +548) spanning the first cleavage site positioned at +414 within the 5’ external transcribed spacer region: 5’-GTGCGTGTCAGGCGTTCT-3’ and 5’-GGGAGAGGAGCAGACGAG-3’ [75]. The human *β-ACTIN* was used as a reference gene and the following primers were used, 5′-TGCGTCTGGACCTGGCTGGC-3′ and 5′-GCCTCAGGGCAGCGGAACC-3′ [76]. The cycling parameters from the manufacturers were followed. 58 °C was used for the annealing step and 42 cycles were performed. Calibration curves based on five serial cDNA amounts (0.016, 0.08, 0.4, 2 and 10 ng/μl) were used for calculation of the reaction efficiencies. The 47S-rRNA expression was normalised to that of *β-ACTIN* according to the C_T_(N^−ΔΔCt^) method, were N represents the primer efficiency, measured in each experiment.

### Lipid Extraction from nuclear fractions

Following cell fractionation, the nuclear pellets were resuspended in nuclear resuspension buffer (10 mM Tris pH 7,4, 1 mM EGTA, 1,5 mM KCL, 5 mM MgCL_2_, 320 mM sucrose) and the number of nuclei counted. Lipids were extracted from each nuclear fraction using a method adapted from Grey *et al* [77]. Nuclei were incubated in 1 mL MeOH/CHCl_3_ 2:1 to extract neutral lipids for 10 min at room temperature with shaking at 1200 rpm and vortexed 3-4 times. The samples were centrifuged at 3000 g for 5 min at 4 °C and supernatants were discarded and the same procedure was repeated. The acidic lipids were then extracted with 750 μL MeOH/CHCl_3_/0.1 M HCl 80:40:1 2:1:0.8 and incubated for 15 min at room temperature and vortexed 4 times during the incubation followed by centrifugation at 3000 g for 5 min at 4 °C. The pellets were resuspended with 250 μL CHCl_3_ and 450 μL 0.1 M HCl and centrifuged at 3000 g for 5 min at 4 °C and a phase split between the organic and aqueous phases was apparent. The organic phase (bottom phase) was collected in conical glass tubes and dried at 60°C under N_2_ gas. Lipids were resuspended with 4-6 μl of MeOH/CHCl3/H_2_O 2:1:0.8, vortexed for 30 seconds before being sonicated in an ice bath for 5 min and vortexed again for 30 seconds. Proteins were recovered from lipid extraction and the protein concentration was estimated for validation of the fractionation by western blotting.

### Lipid Overlay Assay

Lipids obtained from lipid extraction were spotted on Hybond™-CExtra membranes, 2 μl at a time. The membranes were left to dry for 1 hour at room temperature protected from light. The membranes were next blocked for 1 h at room temperature with the appropriate blocking buffer (1% fat-free milk in PBS pH 7.4) and further incubated with 0.5 μg/mL GST-GRP1-PH in the same buffer overnight at 4°C and protected from light. GST-GRP1-PH was expressed and purified as described previously [78]. The membranes were washed 6 × 5 min in PBS-T (0.05% Tween 20) and then incubated with anti-GST conjugated to HRP (1:30 000) in blocking buffer for 1 h at room temperature. The blots were washed 6 x 5 min with PBS-T. The signal was detected by ECL using the SuperSignal West Pico Chemiluminescent Substrate or with SuperSignal West Femto Maximum Sensitivity Substrate (Thermo Fisher scientific) and detected with a BioRad ChemiDoc™ Xrs+. Lipid spot densitometry was quantified using ImageJ.

### Immunostaining and microscopy

Cells grown on coverslips were fixed with 3.7% paraformaldehyde/PBS for 10 min, washed thrice with PBS, permeabilised with 0.25% Triton X-100/PBS for 10 min, blocked for 1 h with blocking buffer (3% BSA in 0.05% Triton X-100/PBS) and incubated with primary antibodies diluted in blocking buffer overnight at 4°C and subsequently with fluorescently-labelled secondary antibodies diluted in blocking buffer for 1 h at room temperature. For labelling with EGFP-GRP1-PH, cells were blocked in 3% fatty-acid free BSA and 0.05% Triton-X100 in PBS for 1 h at RT followed by incubation with 40 μg/ml of the probe in 1% fatty-acid free BSA and 0.05% Triton-X100 in PBS for 2 h at RT. Washes were performed with 0.05% Tween-20/PBS after antibody incubation. The coverslips were mounted in ProLong Gold Antifade Reagent containing 4’,6-diamidino-2-phenylindole (DAPI) or without DAPI and DNA labelling was performed using Hoechst 33342. Images were acquired with a Leica DMI6000B fluorescence microscope using x40 or x100 objectives or Leica TCS SP5 confocal laser scanning microscope using a 63x/1.4 oil immersion lens. Images were processed with a Leica application suite V 4.0 and Adobe Photoshop CC 2018.

### Patient series and immunohistochemistry

Tissue was collected from patients diagnosed with endometrial cancer at Haukeland University hospital during the period from 2001-2013 and included a total of 725 primary tumours (should we add more on the type/grade of the samples here?). Clinical data were collected as described earlier [79, 80]. The patient cohort used for p110β immunohistochemistry is described in detail in Tangen et al [80]. This study was conducted in line with Norwegian legislation and international demands for ethical review and was approved by the Norwegian Data Inspectorate, Norwegian Social Sciences Data Services and the Western Regional Committee for Medical and Health Research Ethics (REK 2009/2315, REK 2014/1907). Patients signed an informed consent. TMA sections were stained and scored for p110β expression following a protocol previously described [80]. Briefly, three cylinders of 0.6 mm were retrieved from high tumour purity areas using a custom-made precision instrument (Beecher Instruments, Silver Spring, MD, USA) and mounted in a paraffin block. TMA sections (5 μm) were stained for p110β expression and scored visually by light microscopy using 20x objective by two independent observers (CK and ILT). Scoring was performed blinded for information regarding clinical characteristics and outcome for the cytoplasmic and the nuclear areas. A semi quantitative and subjective scoring method was used, and a staining index was calculated as a product of the staining intensity score (0, no staining; 1, weak; 2, moderate; and 3, strong) and the area of positive tumour cells score (1≤10%, 2=10-50% and 3≥50%), leading to scores ranging from 0 to 9. The ratio between the nuclear and the cytoplasmic scores was then calculated.

### Statistical analysis

For clinical samples, statistical analyzes were performed using the software package SPSS 22 (SPSS Inc, Chicago, IL) and the values of *P*<0.05 were considered statistically significant. Correlations between groups were evaluated using the Mann-Whitney U test for continuous variables.

## Acknowledgments

We thank Pamela Pollock (Queensland University of Technology, Brisbane, Australia) for providing the EM cells. This work was funded by the University of Bergen, the Norwegian cancer society (project number 2183087 to AEL and TK) and the Meltzer foundation (to FMG).

## Author contribution

AEL, TK and FMG designed the experiments. TK, FMG, APM and VSA carried out most of the experiments and assisted in the analysis and interpretation of data. CK and ILT acquired and analysed immunohistochemistry data. DCT assisted in the acquisition of data. AEL and FMG wrote, extensively reviewed and revised the manuscript. AEL supervised the research.

