## Supplementary table S1 and Figure S1-5 for "Nuclear upregulation of PI3K p110β correlates with increased rRNA transcription in endometrial cancer cells"

**Supplementary Figures S1-S5.**

**Supplementary Table S1.** Antibodies used for immunofluorescence (IMF), Immunohistochemistry (IHC) or Western immunoblotting (WB)

| Antibody | Catalog number | Company | Dilution |
|---|---|---|---|
| **p-S473-AKT** | 9271 | Cell signaling Technology | **WB:** 1:1000 |
| **Total AKT** | 2920 | Cell signaling Technology | **WB:** 1:2000 |
| **Calnexin** | ab22595 | Abcam | **WB:** 1:2000 |
| **GST-HRP** | ab3416 | Abcam | **WB**: 1:30000 |
| **Lamin A/C** | sc-376248 | Santa Cruz Biotechnology | **WB**: 1:10000 |
| **Nucleolin** | 12247 | Cell signaling Technology | **IMF:** 1:100 |
| **Nucleophosmin** | 32-5200 | Zymed/Life Tech | **IMF:** 1:1000 |
| **PtdIns(3,4,5)$P_3$** | Z-P345b | Echelon | **IMF:** 1:400 |
| **PI3K p85α** | 05-212 | Millipore | **WB:** 1:2000 |
| **PI3K p85β** | S3089 | Epitomics | **WB:** 1:5000 |
| **PI3K p110α** | 1683-1<br>4249 | Epitomics<br>Cell signaling Technology | **WB:** 1:5000 |
| **PI3K p110β** | ab151549<br>3011 | Abcam<br>Cell signaling Technology | **IHC/IMF:** 1:50<br>**WB:**1:1000 |
| **α Tubulin** | T5168 | Sigma | **WB:** 1:20000 |
| **Goat anti-Mouse IgG Alexa Fluor 594** | A-11005 | Thermo Fisher Scientific | **IMF**: 1:200 |
| **Goat anti-Rabbit Alexa Fluor 488** | A-11008 | Thermo Fisher Scientific | **IMF**: 1:200 |

### Supplementary Figure 1

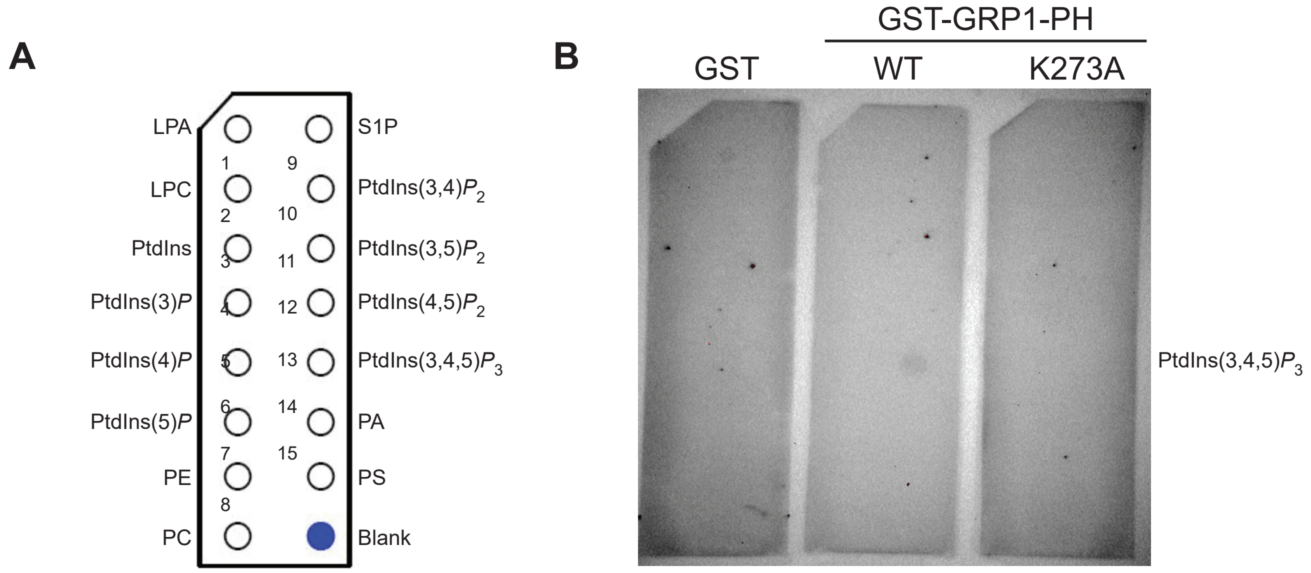


**Supplementary Figure 1. Specificity of PPIn interaction of GST-GRP1-PH.**
**A)** Diagram of PIP strips (Echelon Biosciences) indicating the location of all lipids. LPA, lysophosphatidic acid; LPC, lysophosphatidylcholine; PI, phosphatidylinositol; PE, phosphatidylethanolamine; PC, phosphatidylcholine; S1P, sphingosine-1-phosphate; PA, phosphatidic acid; PS, phosphatidylserine. **B)** Validation of the specificity of the recombinant GST-GRP1-PH WT versus binding mutant K273A by lipid overlay assay.

#### Supplementary Figure S2

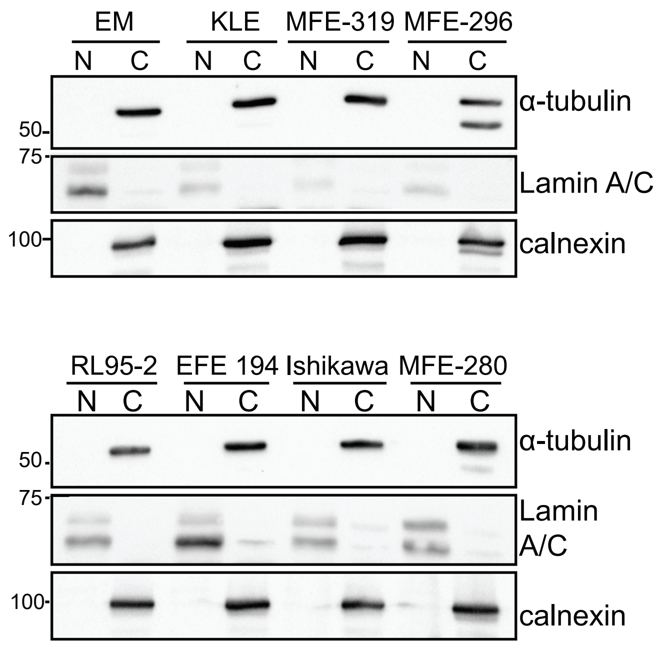


**Supplementary Figure S2: Purity of fractionation**
Actively growing cells were fractionated into cytoplasmic and nuclear fractions. Equal protein concentrations were resolved by SDS-PAGE and analysed by Western immunoblotting using the antibodies as indicated.

### Supplementary Figure S3

**MEF-319**

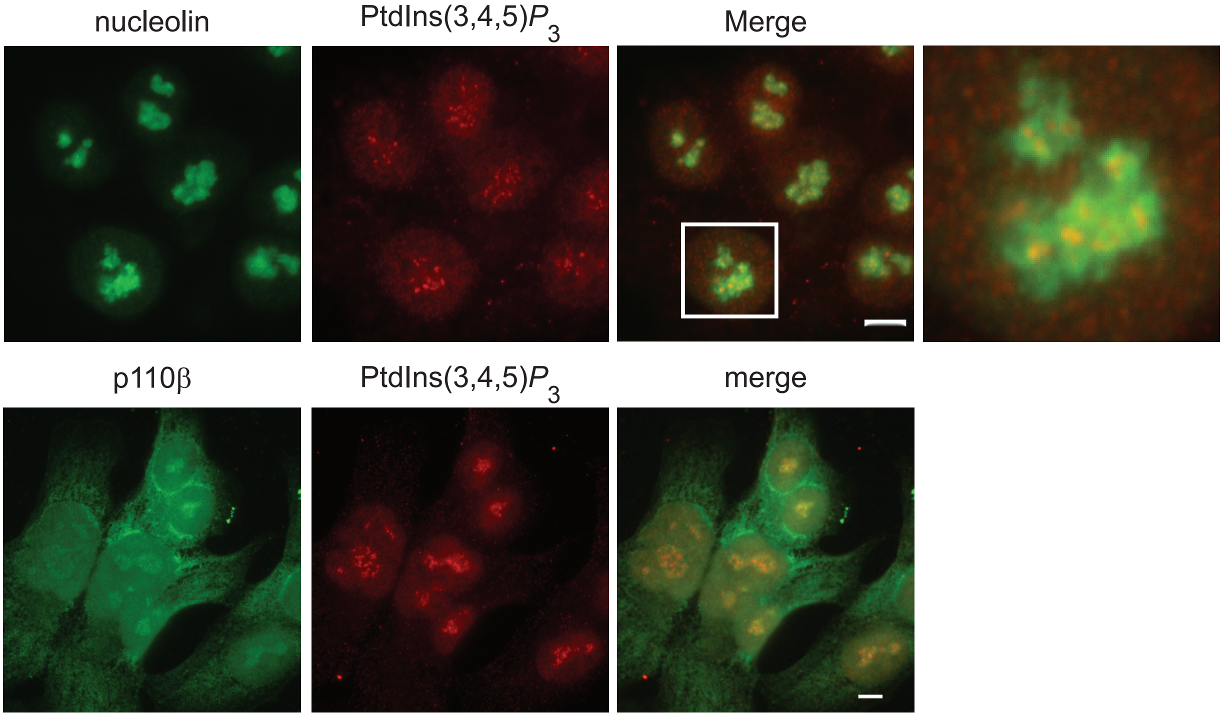


**Supplementary Figure S3: p110β and PtdIns(3,4,5)$P_3$ are nucleolar in MFE-319 cells**
Actively growing MFE-319 cells were immunostained using the antibodies as indicated and imaged using epifluorescent microscopy. Scale bars are of 5 μM.

### Supplementary Figure S4

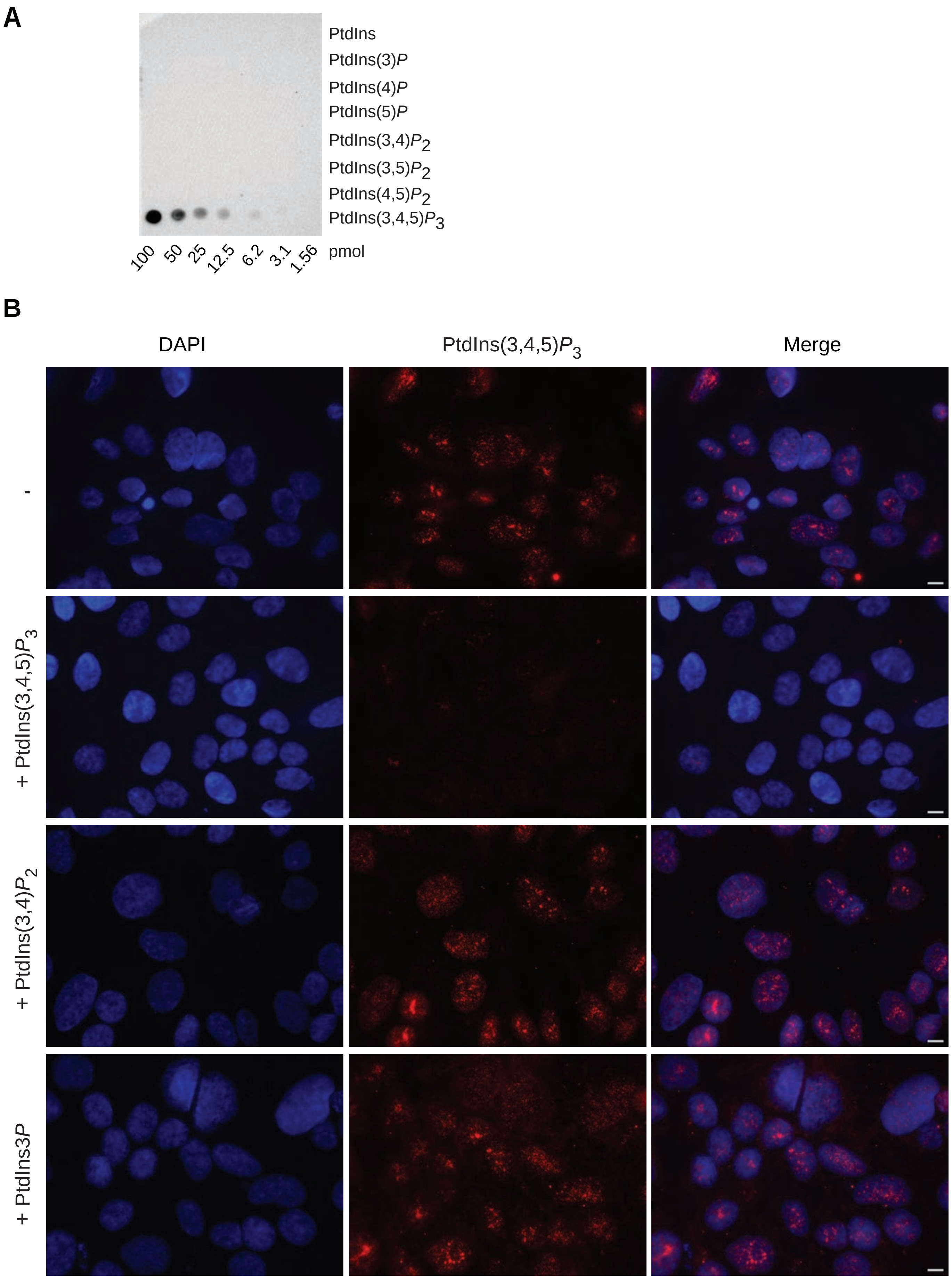


**Supplementary Figure S4: Specificity of the anti- PtdIns(3,4,5)*P*3 antibody**
MFE-319 cells stained with DAPI and an anti-PtdIns(3,4,5)P3 antibody which was pre-incubated without (-) or with (+) 1 μM diC8 PtdIns(3,4,5)*P*3, PtdIns3*P* or PtdIns(3,4)*P*2 and imaged by epifluorescence microscopy. Scale bars are all 5 μm.

### Supplementary Figure S5

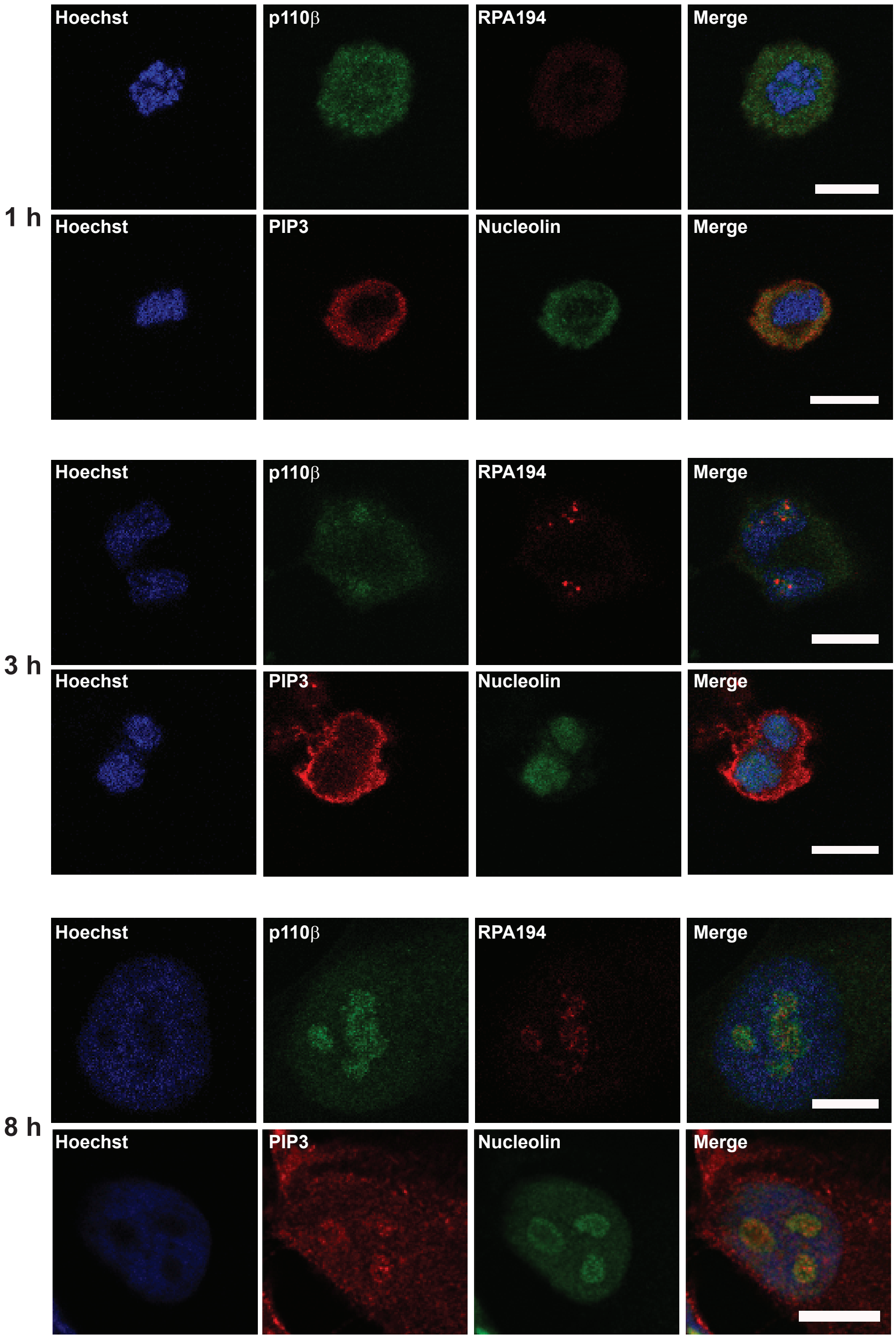


**Supplementary Figure S5**
HeLa were treated for 16 h with 50 ng/ml nocodazole, collected after mitotic shake-off and replated on poly-L-lysine coverslips. Cells were fixed after the different times indicated. Immunofluorescence staining was performed as indicated followed by confocal microscopy. PIP3: phosphatidylinositol(3,4,5)triphosphate, RPA194: RNA polymerase I 194 kDa.
